# Refining sperm quality assessment by high-throughput nuclear morphometric analysis

**DOI:** 10.1101/314708

**Authors:** Anjali A Mandawala, Joe C Perry, Benjamin M Skinner, Grant A Walling, Simon C Harvey, Katie E Harvey

**Affiliations:** Canterbury Christ Church University; University of Essex; Topigs Norsvin; University of Greenwich; The Open University

**Keywords:** Artificial insemination, morphology, nucleus, pig, sperm

## Abstract

Artificial insemination (AI) is commonplace in commercial pig breeding, and as such, ensuring sperm sample quality is of utmost importance to avoid reduced farrowing rates and litter sizes. Here, we have used high-throughput nuclear morphometric analysis to compare pig sperm samples categorised as meeting the AI standard (AIS) or not meeting the AI standard (N-AIS). We find that pig sperm nuclei are asymmetric, and that samples contain phenotypic shape abnormalities that show continuous variation. Samples classed as N-AIS have more abnormally shaped sperm than AIS samples, but conventional analysis misses many. The specific phenotypes identified suggest aspects of spermiogenesis that may be disrupted and indicate future avenues for improving pig sperm quality. This method reveals a significant difference in sperm head morphology between AIS and N-AIS pig sperm samples and has the potential to be further developed as a high-throughput tool for sperm head morphology assessment both in the pig breeding industry and in other species.

## Background

Infertility, defined as the failure to achieve a pregnancy after one year of regular unprotected sexual intercourse, will affect around one in six people. Between 20 and 30% of these cases result solely from male factor infertility, with up to 50% of all instances being at least partly the result of male factor infertility [1]. The analysis of semen parameters is an important part of infertility investigations, with semen volume and pH being documented, and sperm parameters such as concentration, progressive motility and morphology all being routinely observed [2]. Many of these factors are determined during spermatogenesis, the complex process by which undifferentiated diploid spermatogonia become highly specialised haploid spermatozoa, with the process involving complicated cellular, proliferative, and developmental phases [3]. Which single sperm parameter is the most important for fertility is a source of conflict in the literature. Some studies, for example, suggest that sperm progressive motility is positively associated with pregnancy rates [4,5], whilst others do not [6]. Similarly, research has indicated that sperm morphology may be correlated with positive pregnancy outcomes [7–9], but others have suggested that this plays less of a role than previously thought [10].

In agriculturally important species such as pigs, cattle, and sheep, semen analysis followed by artificial insemination (AI) is commonplace in breeding programmes [11], with 90% of sows in the world’s primary pork producing countries resulting from AI [12]. As such, an industry-led working code of practice, the AI standard for pigs has been developed to assure standards of AI are received by pork producers in the UK [13]. Generally, the goals of agricultural animal breeding are to maximise the production of meat at a low cost, and to efficiently disseminate the genetics of particularly valuable animals. As such, the use of high-quality semen doses is of paramount importance, for example in pigs the use of lower-quality doses is associated with lower farrowing rates and smaller litter sizes [14,15]. The fast identification of animals that consistently produce N-AIS samples is therefore important to ensure their removal from breeding programmes; it has been shown for example that boars with fertility problems in a nucleus herd have the potential to reduce litter sizes throughout the entire breeding population [16], in part due to one boar ejaculate commonly being used to serve between 15 and 25 sows [17] in routine AI. Semen parameters are therefore of key importance both in human fertility assessment and treatment and in agricultural practice.

As is the case for all mammals, an ejaculate from a boar, bull or ram does not contain a homogenous population of sperm [18] making analysis and the subsequent identification of subfertile individuals, challenging. In the general analysis of boar sperm morphology for example, a sample is deemed to be “fertile” if the frequency of abnormal sperm heads does not exceed 10%. Fertility can also be assumed if the frequency of abnormalities in acrosomes, mid-pieces, tails, or proximal cytoplasmic droplets is less than either 5% each, or 15% when combined [19]. In humans, the percentage of normal forms (in both fertile and subfertile men) is usually far less than 30% [20], and has been documented to be consistently (and staggeringly) low, with only 3-5% normal forms being identified in assisted reproductive technology (ART) cases [9,21,22].

There also remains much to learn about mammalian sperm biology. Only a limited number of studies to date have focused on the analysis of sperm nuclear morphometry in agricultural animals [23,23,24], and whilst it has previously been shown that sperm shape differs between high and low fertility bulls [25], such observations are limited in pigs [26,27] and differences in morphometric variables that may be related to pig fertility are yet to be elucidated [28]. Whilst the goals of the livestock industry are in stark contrast to human fertility treatment, furthering our understanding of the biology of sperm in general has potential benefits for both fields. Here, we have used Nuclear Morphology Analysis (NMA) [29–33], morphometrics software for microscopy images, to measure and compare the shapes of sperm nuclei from 21,002 pig sperm cells. This is the largest reported number of individual sperm nuclei that have been analysed at this level of detail in any agricultural animal. Our objective is to understand how sperm shape relates to samples being classified as AIS or N-AIS, and to determine if certain sperm features can be correlated with specific biological processes. If these differences can subsequently be linked to fertility, such analysis has the potential to provide an alternative to the livestock industry standard practices and may provide transferrable insights into human sperm morphology.

## Methods

### Semen collection

Fresh ejaculated sperm samples from boars were collected using the gloved hand method [34], by trained staff at JSR Genetics Ltd. Samples were stored in Duragen extender, supplemented with no less than: 500 IU per ml streptomycin; 500 IU per ml penicillin; 150 mg per ml lincomycin; and 300 mg per ml spectinomycin, diluted to 2.3 billion sperm per dose. Samples were stored at 17°C and were prepared within two days of collection. No specific ethical approval was required for this study as all semen doses used were collected as part of JSR Genetics Ltd.’s standard commercial procedures. Semen samples identified as N-AIS by the industry were discarded, and those that were identified as AIS by the industry were used commercially.

### Sample preparation

Prior to preparation of samples for this study, semen samples were identified as either AIS or N-AIS using computer assisted sperm analysis (CASA) followed by manual assessment at JSR Genetics Ltd. Specifically, samples that had a morphology score of above 70% (obtained from CASA) and a motility score of above 4 (1 to 5, 1 being dead and 5 being excellent progressive forward movement) were graded as AIS and those falling below these criteria were graded as N-AIS.

50 AIS and 44 N-AIS samples were used in this study. 2ml of each semen sample was centrifuged at 300g for 5 minutes at 17°C. The supernatant was discarded, the pellet was resuspended in 1.5ml of fixative solution (100% methanol and 100% acetic acid, added dropwise at a 3:1 ratio) and centrifuged at 300g for 5 minutes at 17°C. The supernatant was discarded, and the pellet was resuspended in 1.5ml of fixative solution. 10μl of each sample was then dropped onto the centre of the surface of a labelled (sample ID, date), steam-warmed slide, immediately followed by 10μl of fixative solution. Subsequently, slides were air-dried for two minutes before one drop of fluorescent DAPI (4’,6-diamidino-2-phenylindole) was added to the centre of the slide. Prepared slides were air-dried in the dark, for at least 20 minutes prior to microscopy.

### Image acquisition

An Olympus IX83 inverted fluorescence microscope equipped with a XM10 monochrome CCD camera and CellSens Dimension version 1.9 (expandable imaging software for Life Science microscopy) was used for image capturing. A minimum of 200 nuclei per sample were imaged at 1000x magnification.

### Data analysis

Images were analysed using Nuclear Morphology Analysis (NMA) v2.2.1 [30,33], available here). This software enables automated recognition of round or asymmetric nuclei within an image of interest, and the subsequent morphological analysis of these nuclei. We adapted the landmark recognition to detect and analyse pig sperm nuclei. The software generates ‘profiles’ describing the interior angles around the perimeter of the nucleus, and the diameter and radii across the centre of mass (Figure 1). Pig sperm are (mostly) symmetrical about the anterior-posterior axis, and the tail attachment region, characterised by a ‘dimple’ in the nucleus, was chosen to anchor the profiles and orient the nuclei (Figure 1A, reference point 1).

**Figure 1.**
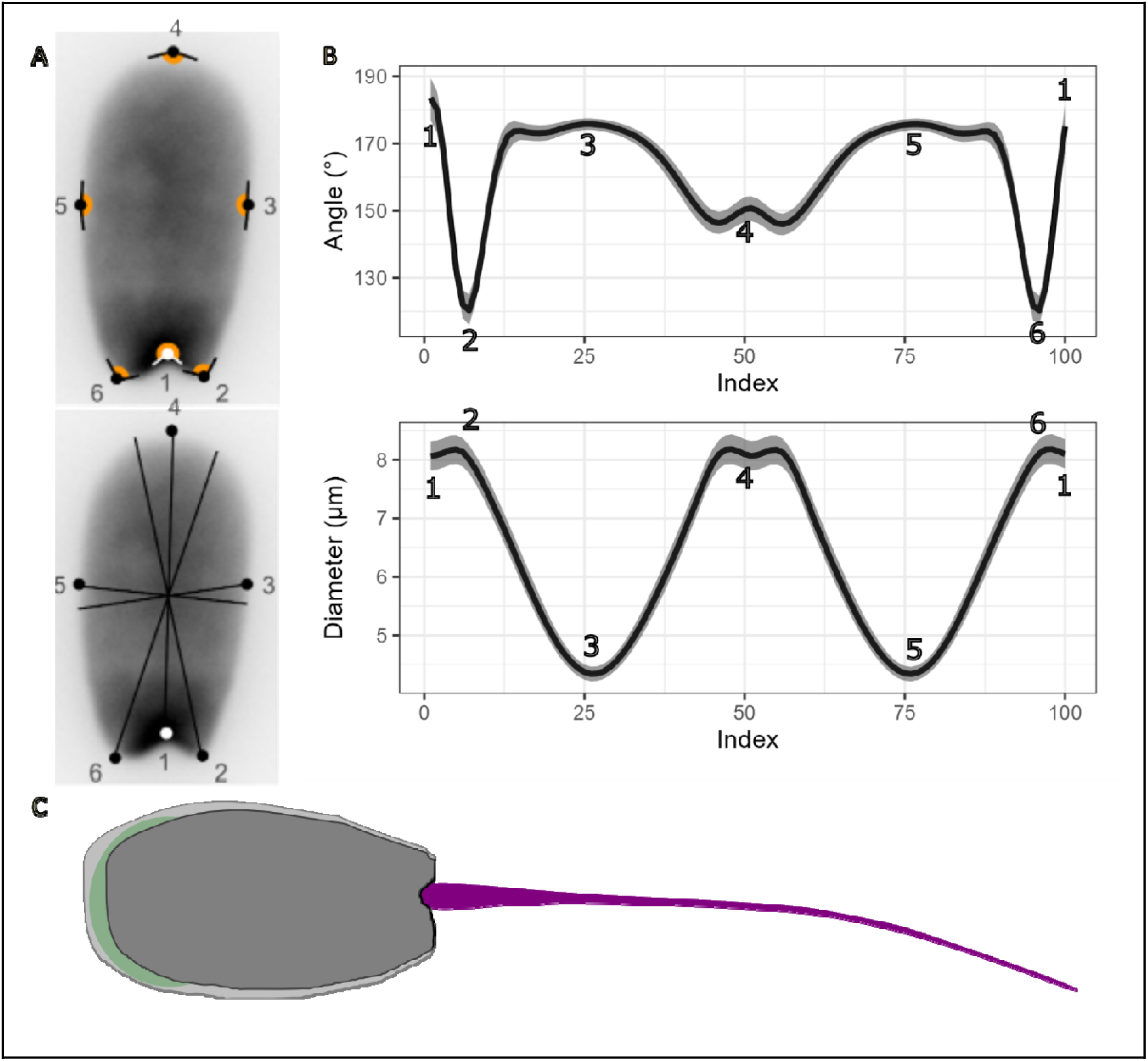
Example morphometric profiles of pig spermatozoa. A) angle profiles measure the internal angle around the perimeter of the nucleus. Diameter profiles measure diameter across the centre of mass. The numbered points are indicated in the profiles, demonstrating key landmarks can be identified. B) Profiles show median and interquartile range of the angles and diameters of cells. C) schematic of a pig spermatozoon; the sperm nucleus (dark grey) fills most of the sperm head, with the remainder of the head comprising the acrosome (green) and the residual cytoplasm (lighter grey). The tail (purple) is not shown to scale but is included to illustrate the attachment region.

For consistent alignment of the nuclei, the tail attachment region (as can be seen in Figure 1 A point 1 and 1C) was placed directly below the centre of mass of the nucleus.

Data was exported from NMA and further processed in R (4.4.0) [35] using a custom analysis pipeline. Initially, k-means clustering on diameter profiles was used to identify distinct groups of shapes, with dimensionality reduction using UMAP [36,37] employed to visualise the shape profiles. Continuous phenotypes were then detected by selective focus on outlier cells, detected using the angle, radius and diameter profiles (Figure 1B).

Cells were classed as normal or abnormal using the profile data. Each profile contains 100 indexes (Figure 1A and 1B); at each index, potentially abnormal cells were detected as those cells with values outside 1.5x the interquartile range. Cells were then scored on how many indexes met the criteria for abnormality. Consistent abnormal phenotypes have higher values, while more normal phenotypes have lower values. For each profile, the cells with a score higher than the mean number of abnormal indexes were determined to be morphological outliers. This allowed us to identify continuous morphological variation across cell populations. Cells were assigned a grouping of ‘normal’ (not identified as outliers), ‘intermediate abnormality’ (the cell was an outlier across a single profile type) or ‘extreme abnormality’ (the was an outlier across multiple profile types).

Associations of breed, sample and date of collection with the severity of phenotype were tested using a Scheirer-Ray-Hare test (a non-parametric equivalent to a two-factor ANOVA). A significant association prompted the use of ordinal logistic regression to explore the association further. The samples were filtered to only known breeds and collection dates for which all samples had data (9321 cells). Ordinal logistic regression was implemented with the *polr* function in MASS [38] using a negative log-log link to account for higher proportions of normal phenotypes, modelling the phenotype cluster as the interaction of collection date and sample. The proportional odds assumption of the model was tested with the Brant-Wald test, and an intercept-only model was created to test the null hypothesis that the predictor variables added no information.

The predictive ability of morphometric data to classify samples as either AIS or N-AIS was tested using logistic regression via generalised linear models. Two models were created; the first using standard industry measurements (width, perimeter, area, ellipticity, elongation, roughness, regularity and height [19,39]), and the second using the morphometric profiles (angle, radius and diameter), modelling a sperm class as dependent on the morphometric parameters. The samples were split into training (70%) and test (30%) groups. The probability of being in the AIS group was calculated for each sperm in a sample, and the mean probability was taken as the overall sample value.

Figures were generated using ggplot2 [40],UMAPs were generated using the UMAP package [36,37], Factoextra was used for heirarchicdal clustering and Caret was used for GLM validation. Scripts used in this analysis are available here.

## Results

### Pig sperm samples contain continuous phenotypic abnormalities

Analysis of sperm nuclear morphology from 50 AIS ejaculates and 44 N-AIS ejaculates yielded measures from 11,416 and 9,586 nuclei, respectively. Comparisons between these groups of animals indicated that sperm heads differ, but that there is a large amount of overlap and variation between individuals (Figures S1-4).

To identify different nuclear shape phenotypes in the samples, all samples were combined into a single dataset to enable a full unbiased comparison of shapes. The first global observation indicated that sperm fell clearly into two equal sizes groups with distinct shape profiles (Figure 2A).

**Figure 2.**
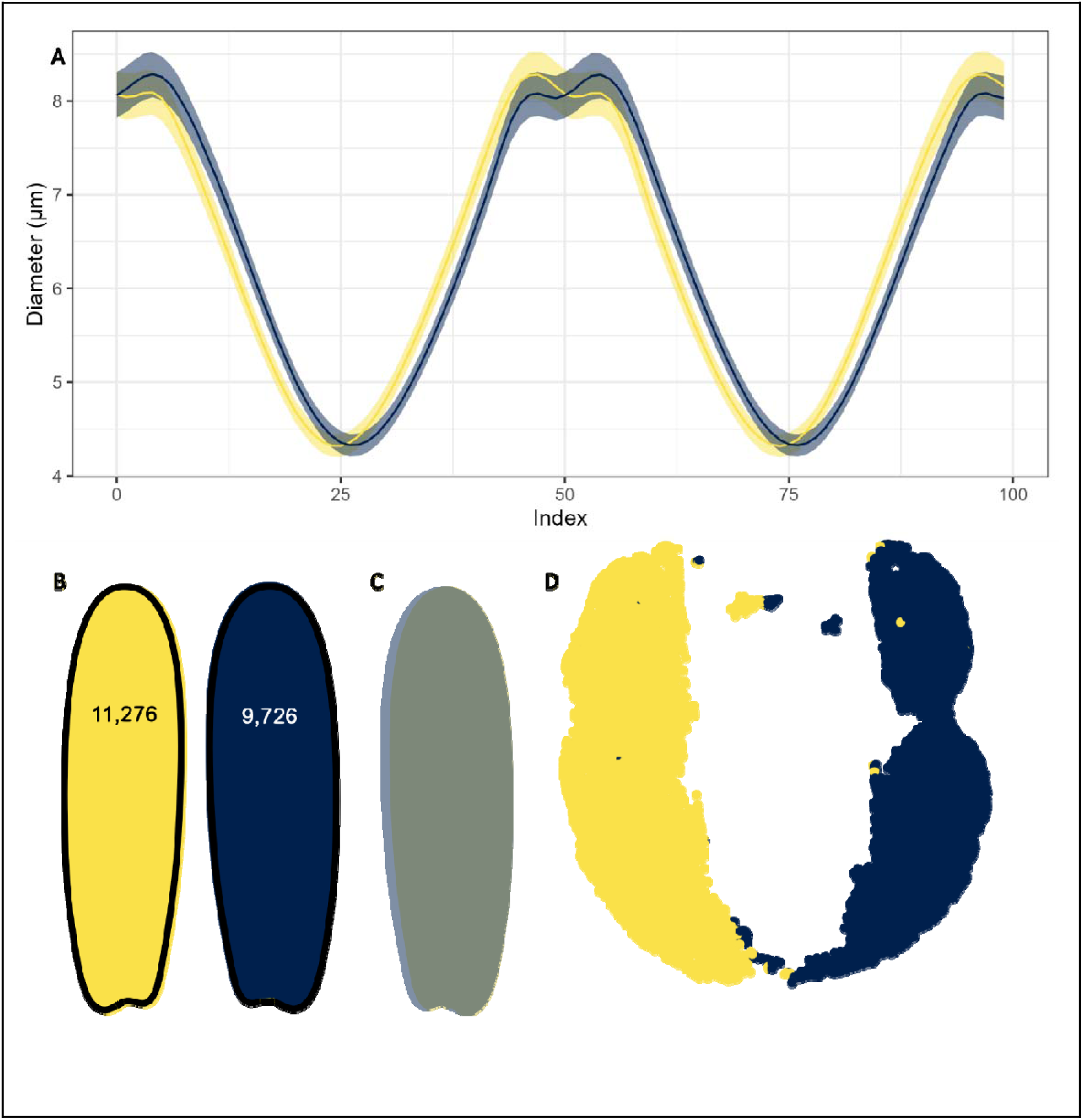
Cell shapes fall into two equally sized groups with mirroring around the vertical axis; A) diameter profiles for each cluster show the symmetrical clusters B) Consensus nucleus shapes show the main difference is at the tail attachment region with the more prominent side to the left or right respectively. C) the left cell in B mirrored across the vertical axis and overlaid on the right cell shows the two cell populations are mirrors of each other. D) Dimensionality reduction of diameter profiles by UMAP shows almost all cells fall cleanly into these two groups.

These two groups of sperm cells correspond to the differences in the shape of the tail attachment region, which is compressed on either the left or the right side (Figure 2B). The profiles of the nuclei with compression on the right side were reversed to determine if the shapes were mirrored and superimposable. The two groups are indeed mirror images of each other around the tail attachment region (Figure 2C), and almost all cells in the dataset fall neatly into these two major groups (Figure 2D). This, plus the general inflexibility of sperm chromatin, suggests the differences are unlikely to be due to compression from the angle of the tail at the time of fixation. Rather, the nucleus appears truly asymmetric, and the difference we observe is caused by the sperm head falling on its dorsal or ventral surface when the slide was prepared.

Since the shape around the tail attachment region was the primary factor that separated nuclei of different shapes, and UMAP showed continuous variation, standard clustering *e.g.* [41] to identify discrete shape groups was not feasible. Instead, we aggregated outlier nuclei with similar shape profiles to identify continuous phenotypes, iteratively grouping nuclei with the greatest differences to normal profiles. This amalgamation of outlier shapes yielded a normal phenotype group (containing 85.2% of nuclei), an intermediate phenotype group (12.7%) and an extreme phenotype group (2.1%). The progression from intermediate to extreme involved condensation of the basal region and bulbing of the anterior region (Figure 3) with two main pathways; one being the compression with failure to elongate resulting a short pyriform phenotype (Figure 3A;7.75%) and the other being abaxial tail attachment and basal compression resulting in an abaxial pyriform phenotype (Figure 3C;6.25%). The intermediate phenotypes are the hardest to categorise as the vary from with minor tapering at the basal region that is difficult to detect to the relatively well-formed side of the spectrum of pyriform cells. This spectrum makes up the largest portion of the abnormality in pig sperm and is highly variable, containing cells that are either the intermediates of a spectrum of abnormality or the end of more minor morphological changes. There is also within the intermediate phenotypes minor tapering and formation of spikes in the basal region (Figure 1B; 1%).

**Figure 3.**
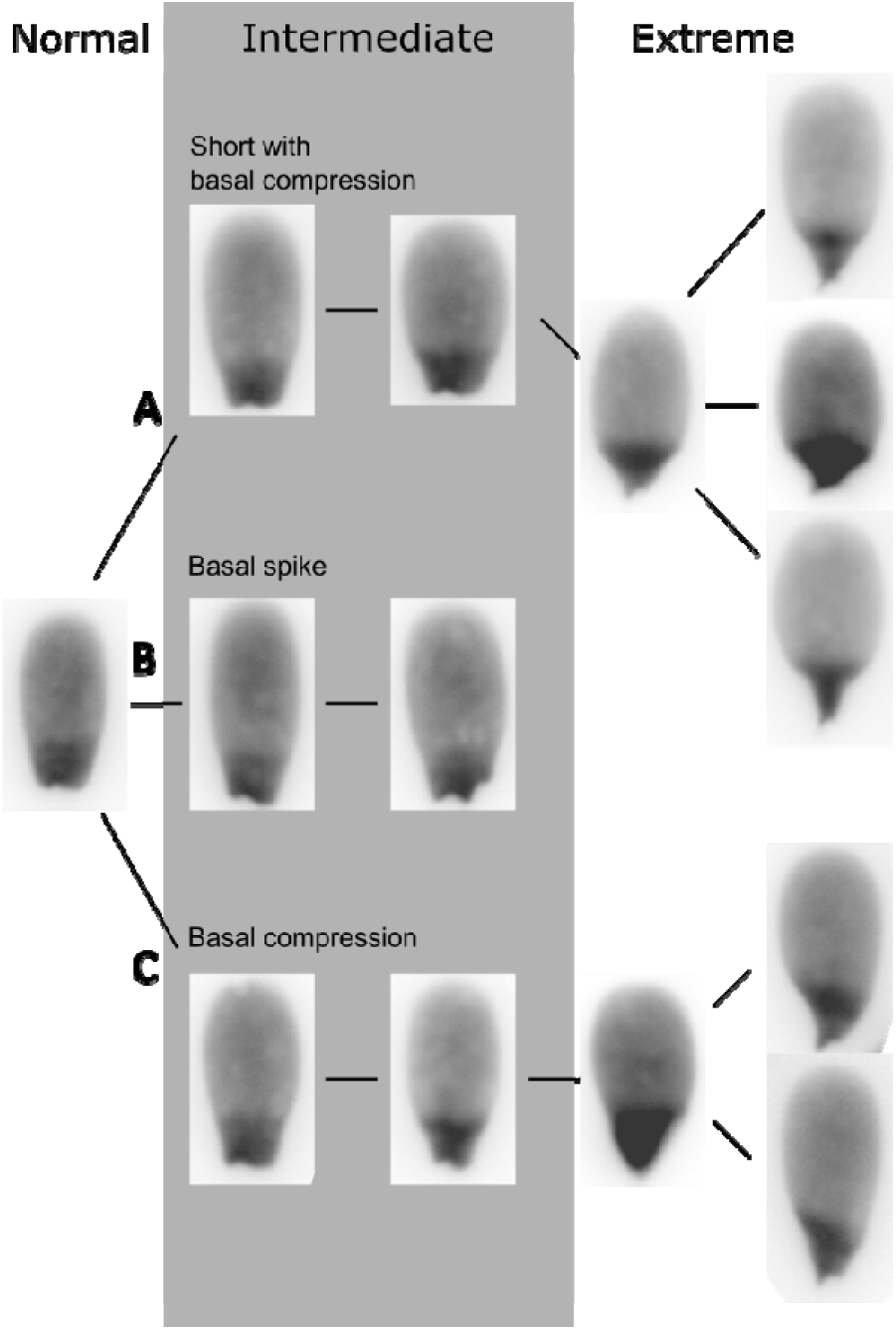
Example DAPI stained images of progressively abnormal sperm phenotypes. Three main classes of abnormality can be seen: A) short (compression anterior-posterior, leading to reduced bounding height) progressing to compression (tapering) around the base of the nucleus and a pyriform phenotype; B) extrusion of nuclear DNA forming a spike near the base of the nucleus with some tapering; C) abaxial compression (tapering) around the base of the nucleus, leading to a pyriform phenotype.

### AIS and N-AIS samples both contain abnormal sperm phenotypes

Our initial analysis indicated that *c.*15% of the sperm were abnormally shaped, but, as samples were pooled, this did not indicate if these were enriched in the N-AIS samples or if their abundance related to other characteristics of the samples. images showing the progressive phenotypes from normal to extreme are shown in Figure 1. We therefore looked at three characteristics: the sample’s status as AIS or N-AIS, the breed of the boar (where known, n= 11,448 sperm heads) and the date of sample collection (as a proxy for environmental or technical factors). Intermediate and extreme sperm phenotypes were not evenly distributed across all samples (Figure 4), and while the abnormal phenotypes were more prevalent (1.96:1 N-AIS:AIS more extreme, 1.44:1 N-AIS:AIS more intermediate) in the N-AIS samples, they were also found in AIS samples.

**Figure 4.**
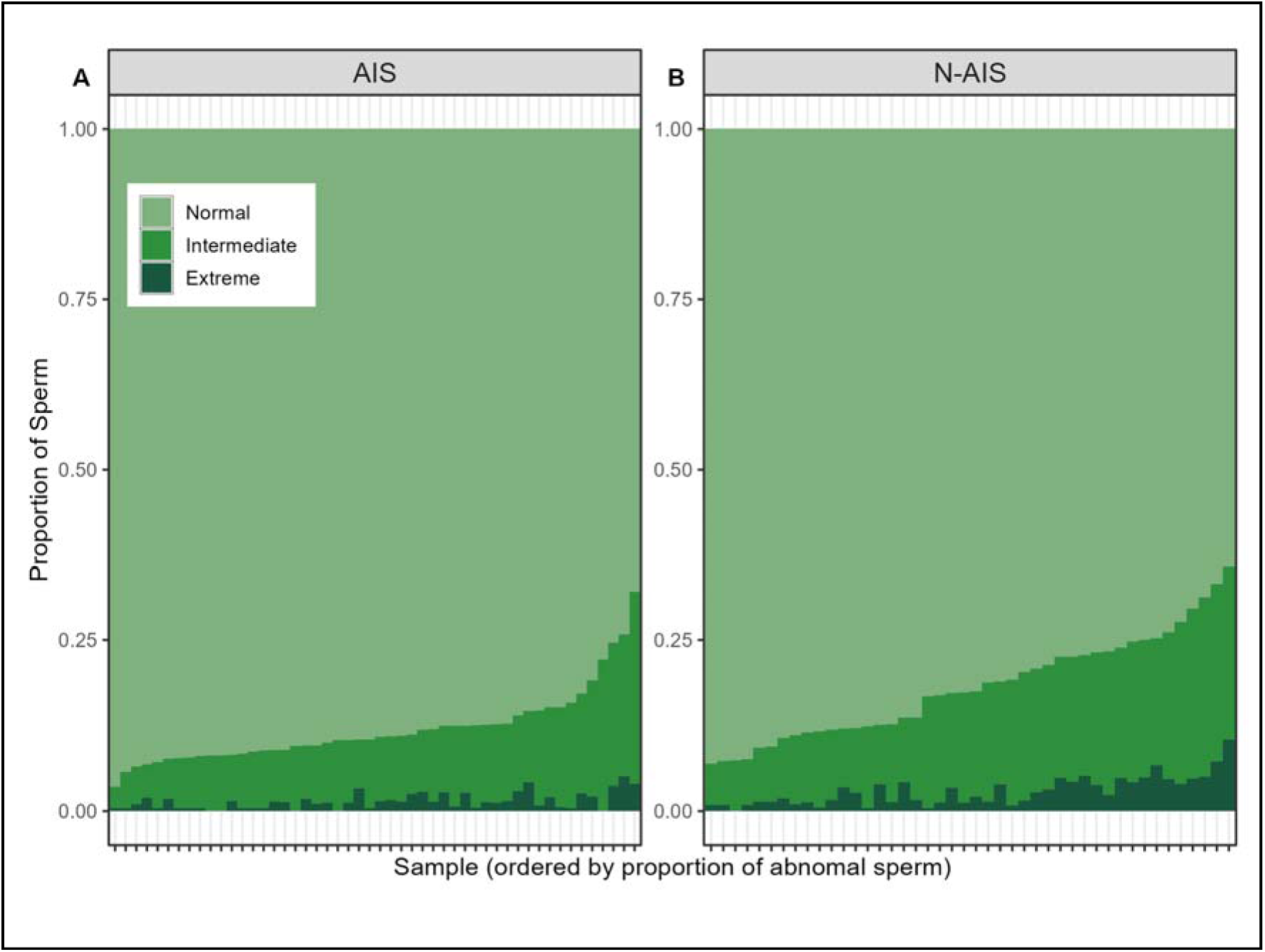
Proportion of sperm in normal, intermediate or extreme groups for A) AIS samples (50 samples) and B) N-AIS samples (44 samples). Intermediate and extreme phenotypes are more frequent in N-AIS samples but are also common in a subset of the AIS samples.

Given that the pattern of data suggested there was variation between samples and between sample collection dates (Figures S3 and S4), we tested whether the distribution of phenotypes could be associated with the date of sample collection or individual samples across the entire dataset. Significant associations were found for both variables and their interaction (Scheirer-Ray-Hare test, p<0.001 for each). To explore the interactions in more detail, we focussed on the samples for which breed was known. We then selected dates for which all these breeds were present. Ordinal logistic regression did not reveal a global association between the severity of the phenotype and the individual sample for the known breeds (p>0.05) apart from White Duroc (p=0.01) in which severe phenotypes were more common (Table S1). The odds that a sperm cell had an extreme phenotype versus intermediate or normal was increased 2.1 times compared to other breeds.

The date of collection had some global associations with severity of phenotype; for example, for samples collected on one date (6^th^ July), the odds that a sperm cell had an extreme phenotype versus intermediate or normal was increased 3.3 times compared to other dates (p<0.001). Most other dates did not show global effects.

Interaction effects were detected between the date of collection and individual samples (see Table S1 for a full breakdown), indicating that the environment has a larger impact on the quality of the ejaculate than the breed of the boar. This was confirmed by testing for differences in the shape profiles between breeds. Repeated Wilcoxon rank-sum tests, multiple testing corrected with the Holm method showed no significant difference in the angle, radius or diameter profiles between breeds suggested that there is subtle nuclear morphological variation between breeds, with differences dominated by shared genetic or environmental factors.

### Classification of normal/abnormal cell types does not align with industry assigned AIS/N-AIS classifications

We next considered if the extra morphometric information measured allowed for more robust classification of samples as AIS/N-AIS than is currently possible. The samples were originally classified based on motility and shape, with 70% morphologically normal forms and a motility score of above 4 leading to a classification of AIS. Although motility information was not available for these samples, the predictive power of conventional morphometrics was compared. For this, a GLM was created to predict the probability of a cell as originating from an AIS or N-AIS sample based on the width, perimeter, area, ellipticity, elongation, roughness, regularity and height, as are commonly used in industry. Each sample was scored as the mean of its individual cell predictions (Figure S2A). This score was then compared to a GLM that used the shape profiles for prediction (Figure S2B). This comparison indicated that greater discrimination between the AIS and N-AIS samples was possible. The ability to classify samples directly by the proportion of normal, intermediate or extreme phenotypes is in principle a useful improvement on the simpler morphometric measures; AIS samples with a high rate of shape abnormalities in particular show that there is morphological variation present that needs to be further explored.

## Discussion

### Continuous sperm shape phenotypes unify previous understanding of pig sperm variation

Studies of pig sperm shape, and indeed the sperm of other species, usually focus on simple morphometric measurements such as area and circularity [19,28,42–44]. However, these measures can miss subtle phenotypes, especially when large numbers of sperm are being screened. Here, 21,002 sperm nuclei have been analysed at high resolution allowing in-depth comparison of sperm nucleus shape. Previous pig sperm morphology studies report the analysis of between 3,000 and 5,550 sperm heads [43,45]. To the best of our knowledge, this is therefore the largest reported number of individual sperm nuclei that have been analysed at this level of detail in any agricultural animal. The use of this large sample size has allowed us to detect and characterise subtle phenotypes that have previously been overlooked.

Previous studies measuring bright-field images have identified differences in sperm head size between pig breeds with Saravia *et al.* concluding that Duroc sperm heads were 0.2μM longer and 0.1μM wider than other tested breeds [19], and Kawarasaki, Enya and Otake finding Duroc sperm heads to be 0.1μM longer and 0.03μM wider [45]. Given that such differences were not observed here (Figure S1), the size difference seen in other studies may be due to the structures outside of the nucleus, or due to sample variation. Of note however, is that the Duroc pig samples analysed in this study were from crossbred white Duroc animals. This crossbreeding may have resulted in the smaller sperm observed due to a lack of genetic diversity or a potential founder effect (Figure S1).

As shown in Figure 2, asymmetry in pig sperm nuclei was identified, with one side of the tail attachment being consistently larger than the other. Given that pig sperm nuclei are flattened, we conject that this asymmetry reflects how the sperm cells fell on the slide during preparation; either dorsally or ventrally. Evidence from scanning electron microscopy also suggests that the asymmetry is present in the full sperm head, not just the nucleus [45].

The other shape phenotypes that were discovered in this study were continuous, with progressive abnormalities involving compression around the base of the nucleus (Figure 3). Some of these shape phenotypes are similar to those reported before, including the division of intermediate and extreme phenotypes [43,45,46].

Basal nucleus compression causing a bulb-like shape has been observed in the sperm of other species such as humans [46], and has been studied extensively in cattle [47–51]. Notably, *Kif3a* knockouts in mice have also been shown to generate similar sperm phenotypes [52], although much of the characterisation of sperm phenotypes to date has been either manual or using CASA systems measuring simple morphometric values, and therefore lack the power required to detect subtle variations [23,24,28,43,45,53,54].

Some of the samples with the greatest proportion of abnormal sperm were collected on the same day, suggesting that environmental factors, either pre- or post- sample collection, may be as, if not more, important than genetic factors, as borne out by previous studies [55–57]. Purely technical factors in sample collection seem implausible to explain the range of phenotypes we observed. Whilst breeding seasons are unlikely to be the cause (as pigs are nonseasonal), it has been shown that stress and physical trauma can affect spermiogenesis, with stress-induced infertility being previously been reported in mice, boars and humans [56,58,59]. Although chromatin integrity is important for fertility [60], morphological differences head shapes are not directly due to chromatin integrity in stud boar sperm [19] or in humans [61] [62] except in the very round sperm examples seen in very few of in pig samples. This suggests that chromatin integrity is not an important factor in the structure of the main phenotypes, as there are very few other sperm that show known oxidative damage-induced phenotypes. It is unlikely that there would be such a disparity in phenotype representation if they were related.

### Progressive pig sperm phenotypes match known defects in mechanisms of spermiogenesis

The sperm were organised into three groups that illustrate the progressive abnormalities (Figure 3). These are (1) extrusion of a basal spike; (2) basal compression, developing asymmetrically; and (3) basal compression with failure of the remainder of the nucleus to elongate. Potential mechanisms that can generate these phenotypes have been described in mice, and information from other mammals can be used as an analogue for boar spermatogenesis [63,64].

The elongation of the nucleus is shaped by five core elements. First, the ectoplasmic specialisation of the Sertoli cell, which provides compressive forces externally [65,66]. Second, the acroplaxome (a cytoskeletal plate composed of f-actin and keratin) upon which the Sertoli cell pushes for nuclear elongation [65,67,68]. Third, the Sertoli cell-spermatid linkers (afadin/nectin complexes) which transmit the forces from the Sertoli cell [69]. Fourth, the manchette (a network of microtubules around the base of the sperm head) which directs the basal nuclear structure and tail attachment [52,70,71]. Fifth, the groove belt (the connection between the machete and the nucleus where the acroplaxome ends; [52,70,71]. These components are interconnected, and while not the only mediators of sperm shape, they are important for understanding how perturbations to the process of elongation in spermiogenesis can cause sperm defects. Many proteins are involved in spermatid elongation; SUN3, SUN4 and SUN5 are of particular importance in the process of manchette formation and the attachment of the sperm head to the tail. Both SUN3 and SUN4 knockouts result in a phenotype that does not successfully elongate and have a round nucleus with a mispositioned acrosome [70,72,73], with SUN3 knockout also causing complete failure of manchette formation [74]. SUN5 anchors sperm heads to their tails [75] and does not affect the acrosomal attachment [76]. SUN5 knockouts form acephalic spermatozoa [75]; these are unlikely to be the cause of the abnormal sperm in our samples as we did not see these phenotypes. Other proteins are more likely candidates to explain these abnormalities. The dynein and kinesin motor proteins are involved in elongation and the action of the Sertoli cell ectoplasmic specialisation. Dynein knockouts in mice result in continuous deformation of sperm with malformation of the general structure with reduced elongation and poor formation of the structures around the acrosome [77–80]. Kinesins are also involved [65] and knock outs of the kinesin *KIF3A* give rise to bulb-like morphology with over compression of the manchette [52].

Nectin-2/3 is part of the spermatid-Sertoli cell connections directing the majority of sperm head elongation [65]. Nectin-2/3 disruption in spermatogenesis affects fertility [81], and is linked to environmental changes [56]. Deficiency of Nectin-2 has also been shown to give a diverse range of abnormal sperm in mice similar to the general structural abnormality shown in the pig sperm [82].

Whilst Figure 3 showed the progression of phenotypes with increasing severity, we outline the structural elements that influence sperm shape in Figure 5 and indicate the forces that may be operating on the developing sperm to generate these phenotypes.

**Figure 5.**
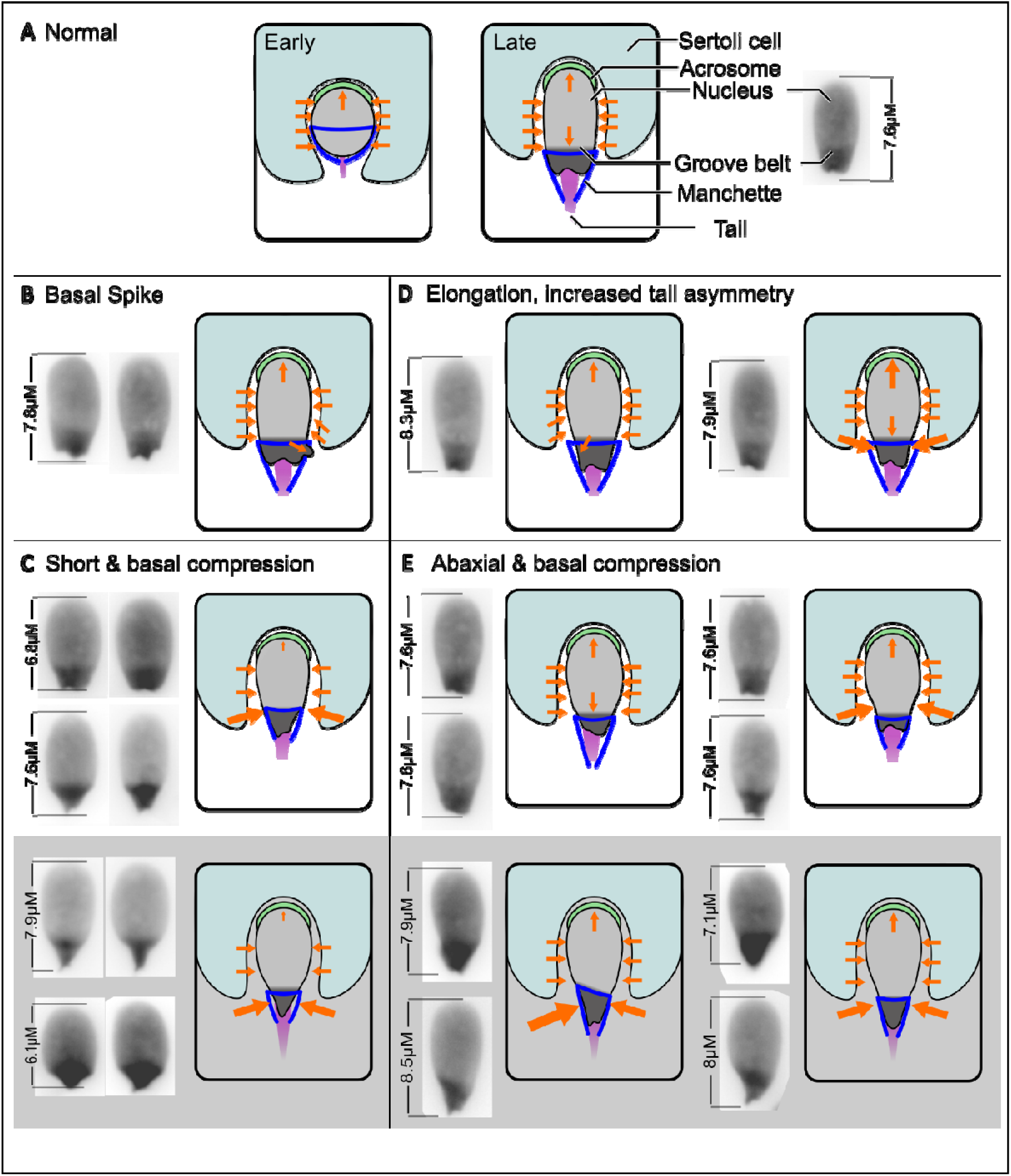
Summary of the forces involved in shaping the sperm nucleus and the abnormal phenotypes we observe. (A) the normal forces of elongation of a pig sperm exerted on and generated within the cell by the Sertoli cell ectoplasmic specialisation. The Basal spike (B) section shows the forces happening to the cells with an anterior bulge around the tail attachment region that is possibly either from the force of a broken tail or malformation of the manchette/ ectoplasmic specialisation hoops slipping due to lack of acroplaxome attachment proteins below the manchette forcing out the bulge. The short with basal compression (C)phenotype has concentrated pressure around the base and a lack of intranuclear pressure which could come either from a lack of molecular motors or misalignment of ectoplasmic specialisation hoops that results in a lot of basal pressure causing constriction but not the elongation seen in other phenotypes. The elongation with increased tail asymmetry (D) is potentially due to manchette misattachment causing the asymmetry that is not severe enough to cause a full abaxial structure or abaxial misalignment of the hoops from the ectoplasmic specialisation without enough constricting force to cause full abaxial development. The elongation of this sperm phenotype is likely due to increased intranuclear pressure from the compression of the manchette and loops that are potentially due to increased molecular motor activity such as dyneins or kinesins or potentially moderated by the same acroplaxome attachment complex that allows for more force transfer due to more attachment. The Abaxial (E) phenotype with basal compression is likely due to a combination of manchette misalignment exacerbated by the basal compression due to the previously mentioned processes.

We see over compression of the manchette and a dysfunction of nuclear elongation; potentially the absence or reduction in levels of one of these connector proteins or molecular motors would result in over-compression of the manchette and improper elongation [52,82]. One potential mechanism for this could involve the over-tightening of the hoops of the Sertoli cell ectoplasmic specialisation via uneven pressure exerted on the nucleus, either because the hoops lack constricting forces or because they lose their solid connection to the acroplaxome and prevent its function as a modulator of elongation.

Although we cannot be certain that these are the mechanisms at play in these boar samples, our analysis indicates that these structural changes are localised to specific regions of the sperm. This suggests that further study of the proteins involved in these regions will be important to understanding variation in boar sperm development. Particularly, if the components are affected by environmental factors, there may be possible interventions to improve sperm quality and ejaculate consistency.

### Implications for the pig breeding industry

This work, and Barquero *et al.* [83] both show a consistent distribution of morphological phenotypes. Barquero *et al.,* however, only used samples comparable to those in our study categorised by breeders as AIS. Given the previously reported link between sperm head size and motility with simple morphological parameters such as height and width [84], and an association between the number of stillbirths in Piétrain boars and poor sperm motility [83], we see clear support for the well-established idea that boar semen quality is significantly affected by the environment [55,57,85]. Further investigation of this relationship is needed, particularly focussing on candidate proteins that can explain the phenotypes; a deeper understanding of sperm morphology may further explain the relationship between morphology, motility and farrowing outcomes.

Evidence suggests that pig AI may be suboptimal [86–89], with this potentially due to the methods used to classify sperm at AIS or N-AIS. This interpretation is supported in this study, where a large overlap of morphology and the subtle intermediate phenotypes of abnormal sperm between AIS and N-AIS samples is seen. It would therefore be an important next step to test the fertility of samples classified as such using this new approach.

## Conclusions

In this study we demonstrate that morphometric analysis of pig sperm using NMA software effectively distinguishes morphologically distinct populations of nuclei. Our analysis reveals differences in sperm nucleus morphology between animals classified as commercially acceptable (AIS) and unacceptable (N-AIS) based on assessment to industry standards. Use of NMA has offered additional insights into pig sperm. Importantly, the distribution of sperm phenotypes between different collection dates and between breeds suggests that environmental factors in sperm collection and transport influence can greatly influence sperm morphology and fertility. Further study is needed to understand how these phenotypes develop, and the functional consequences for boar fertility. These methods may also be of use in the categorisation of the phenotypic distributions within human sperm and have the potential to provide insights into the various complex and subtle morphological differences that could be linked to environmental factors, fertility, and disease.

## Acknowledgments

We would like to thank JSR Genetics Ltd. for supplying the samples that were used in this study.

