## Supplemental Figures S1-S4 for "Refining sperm quality assessment by high-throughput nuclear morphometric analysis"

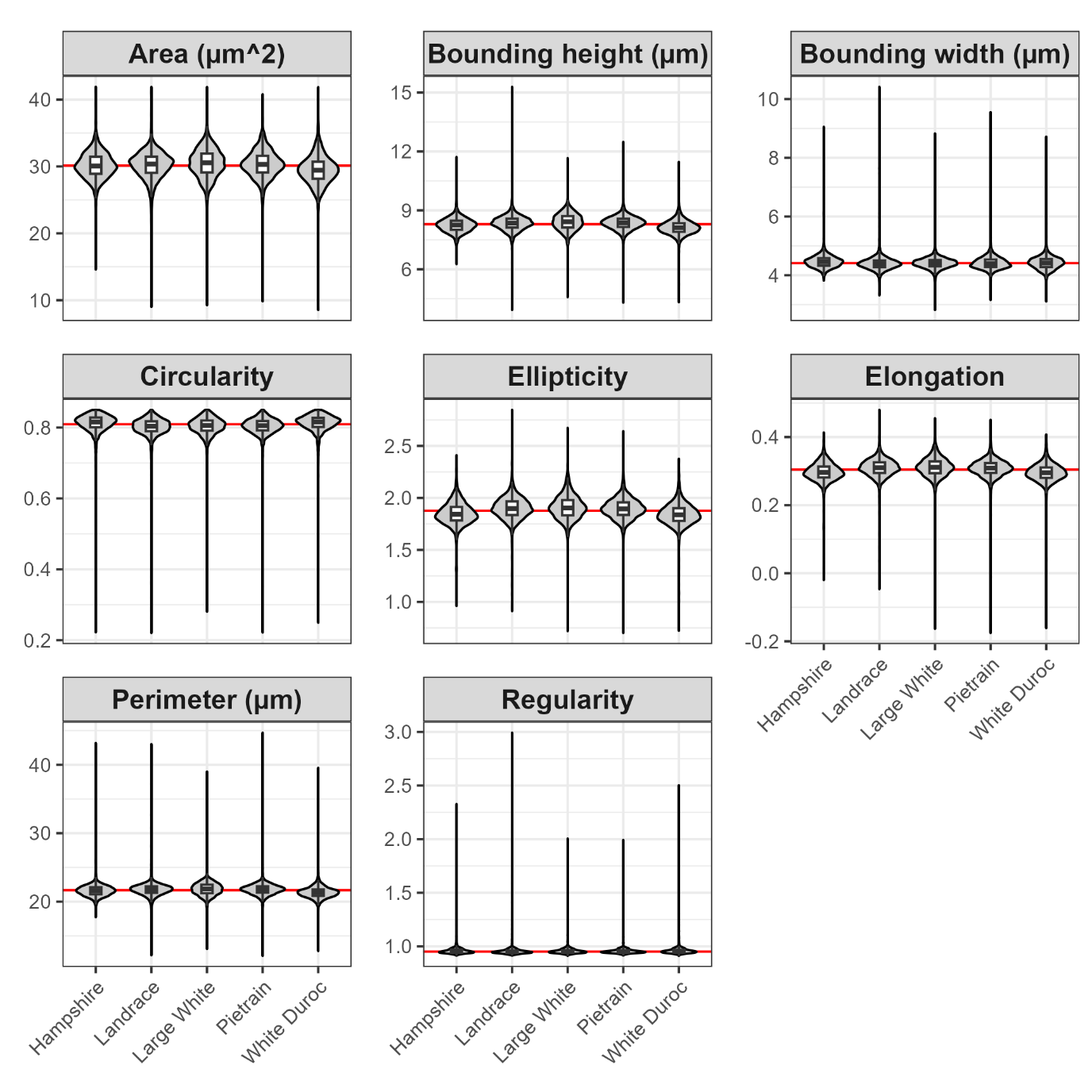

Figure S1: Violin plots showing the industry-relevant sperm cell dimensions: area (µm^2^), bounding height (µm), bounding width (µm), circularity, ellipticity, elongation, perimeter (µm^2^) and regularity, for Hampshire, Landrace, Large White, Piétrain and White Duroc pig breeds.

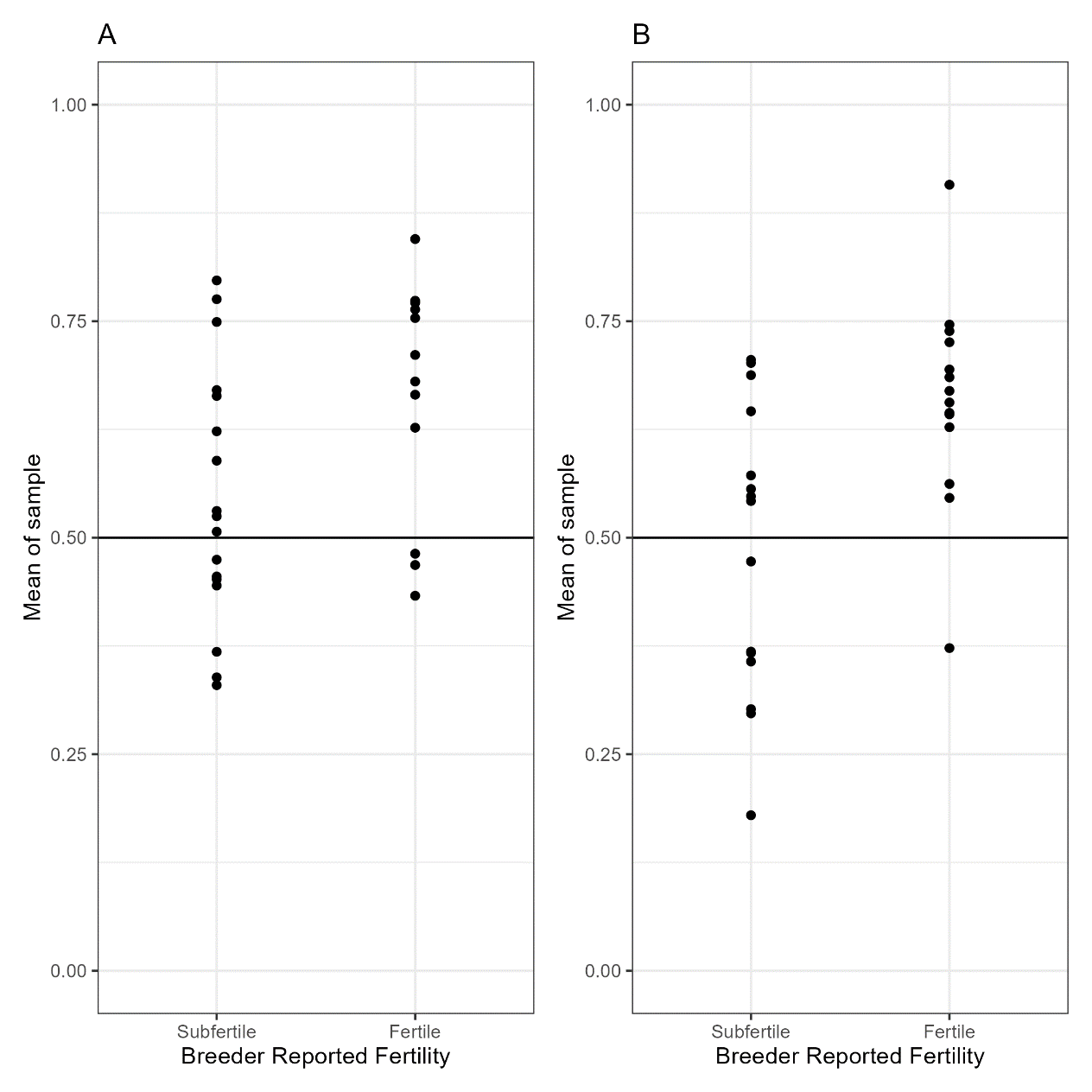

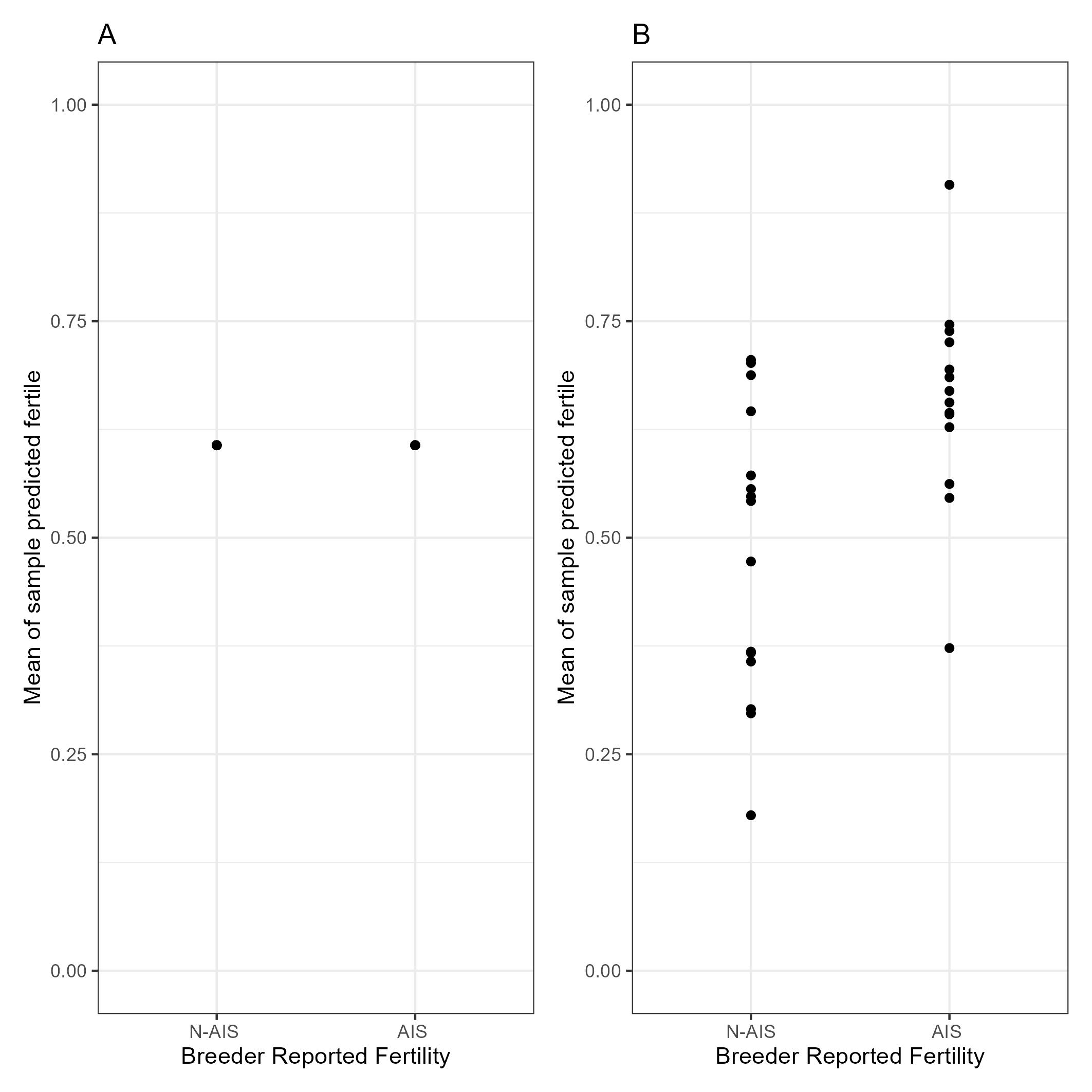

Figure S2: A: N-AIS and AIS samples scored as the mean of individual cell predictions and B: N-AIS and AIS samples compared to a generalised linear model with predictions based on sperm cell shape profiles.

 
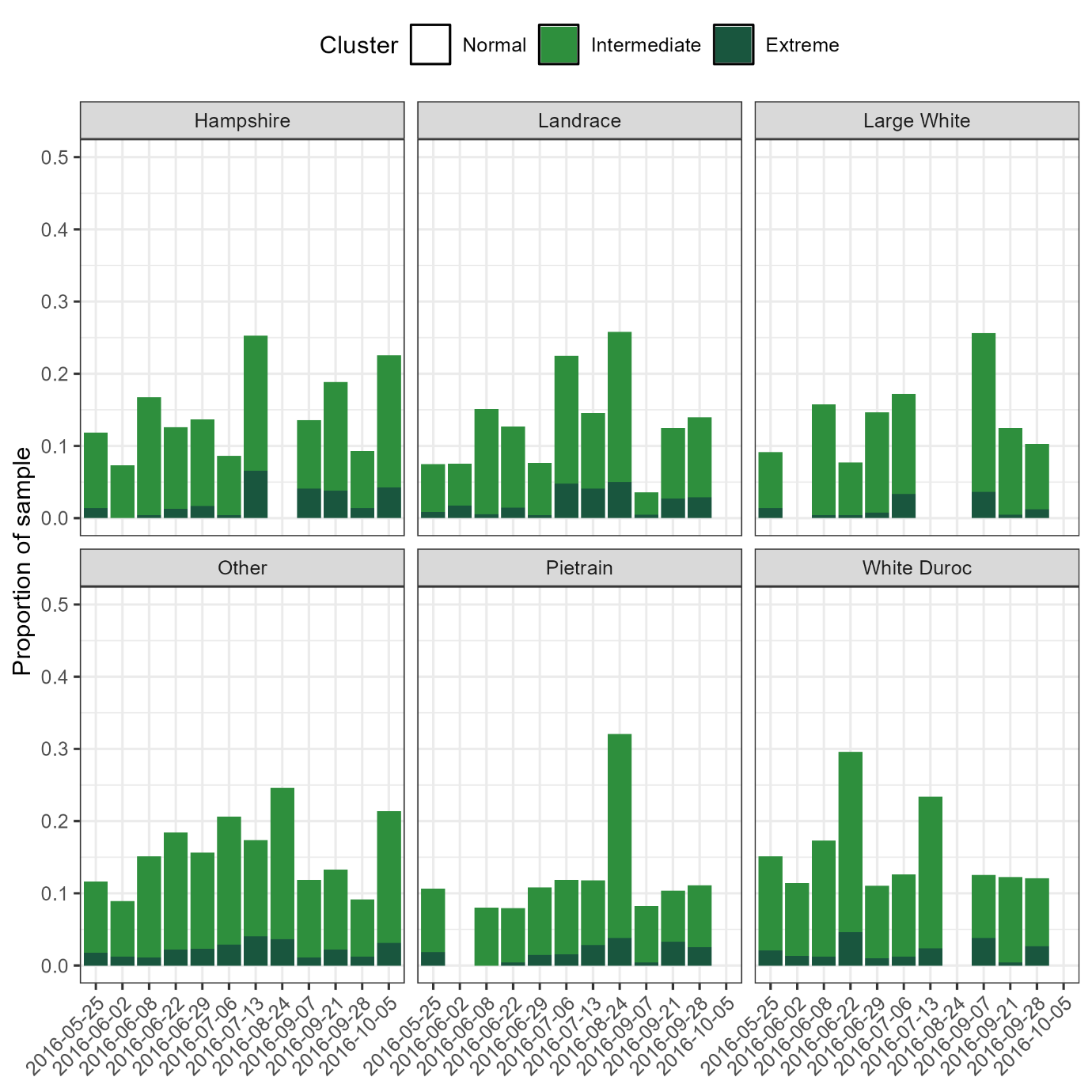

Figure S3: Variation between sperm samples collected on different dates in the proportion of sperm in normal, intermediate or extreme groups for Hampshire, Landrace, Large White, Piétrain and White Duroc pig breeds.

 
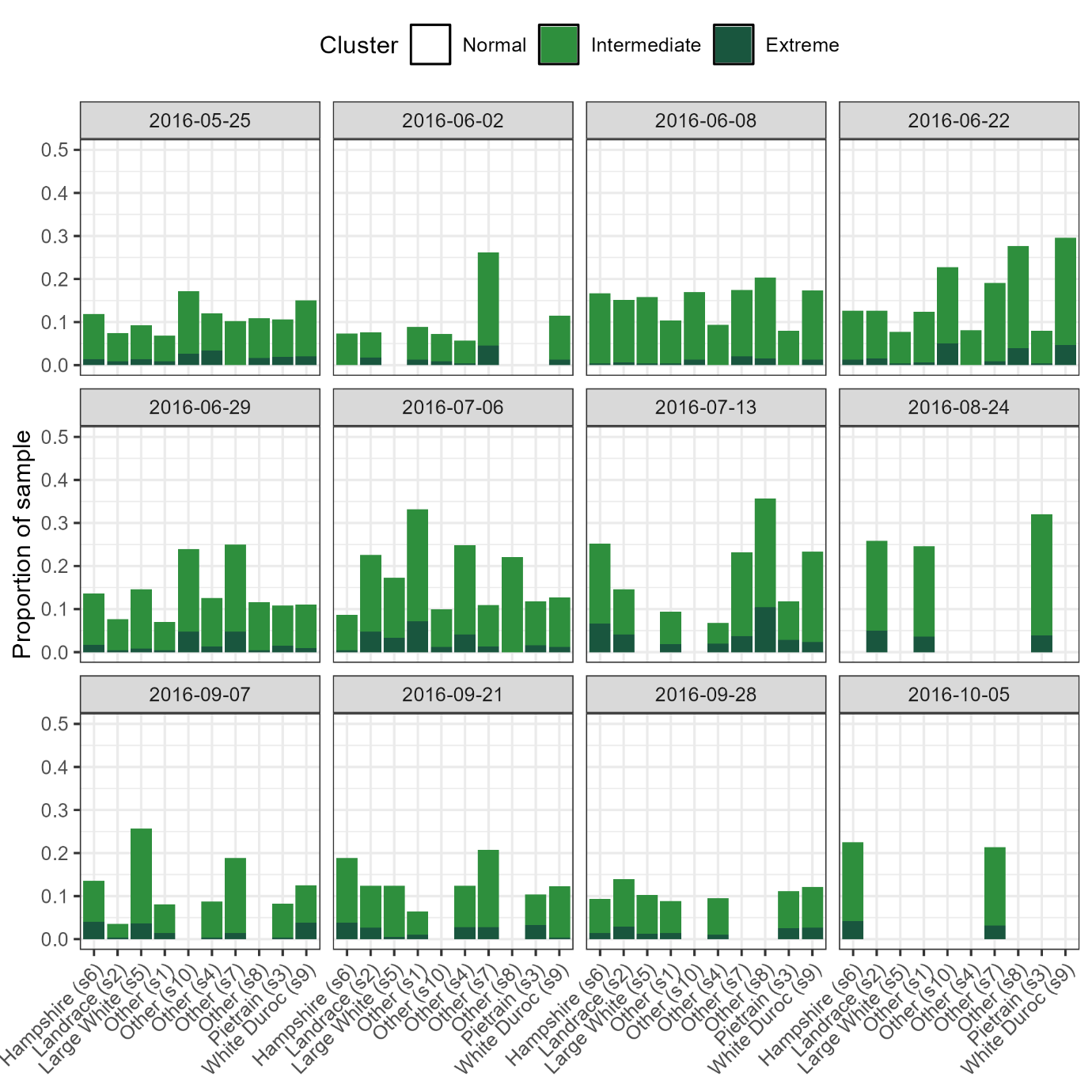

Figure S4: Variation between sperm samples from Hampshire, Landrace, Large White, Piétrain and White Duroc pig breeds in the proportion of sperm in normal, intermediate or extreme groups for samples collected on different dates.
